# CD20^+^ T cells are associated with inflammatory responses in experimental arthritis

**DOI:** 10.1101/2021.10.12.464081

**Authors:** Miguel Pineda, Piaopiao Pan, Yilin Wang, Aneesah Khan, Mukanthu H. Nyirenda

## Abstract

CD20^+^ T cells comprise a small but highly inflammatory subset that has been implicated in autoimmunity, including rheumatoid arthritis (RA). We sought to characterise the CD20^+^ T cell subset at the site of inflammation in murine collagen-induced arthritis (CIA) model of RA and investigate the phenotype and functional relevance of CD3^+^CD20^+^ T cells in the lymph nodes and arthritic joints using flow cytometry and immunohistochemistry. We demonstrate that CD3^+^CD4^+^CD20^+^ and CD3^+^CD8^+^CD20^+^ T cells are expanded in the draining lymph nodes of CIA mice. In addition, compared to naïve mice and those that did not develop clinical symptoms, CD20 expressing T cells of arthritic mice produced increased levels of pro-inflammatory cytokines (GM-CSF, TNF-a, IL-17, and INF-g). Notably, CD3^+^CD4^+^CD20^+^ and CD3^+^CD8^+^CD20^+^ T cells of disease mice were enriched with CXCR5^+^PD-1^+^ T follicular helper cells and CXCR5^-^PD-1^+^ peripheral T helper cells, subsets of T cells that have been implicated in promoting B-cell responses and antibody production within pathologically inflamed non-lymphoid tissues in RA. Importantly, CD3^+^CD20^+^ T cells were detected in the inflamed regions in the lymph nodes and paws of arthritic mice. Our findings suggest that CD20^+^ T cells are associated with inflammatory responses in the arthritic joint and may exacerbate pathology by promoting inflammatory B cell responses.

## Introduction

Autoimmune conditions such as rheumatoid arthritis (RA) are characterized by lymphocytic tissue infiltrates containing aberrant T and B cells and innate immune cell populations that play a critical role in disease pathogenesis (1). Various subtypes of T cells, including Th1 and Th17, take part in immune-mediated inflammation of RA, where they become activated and accumulate in the inflamed joints (1, 2). These T cells produce pro-inflammatory cytokines (3, 4), and help B cells proliferate and secrete essential proteins such as rheumatoid factors (RFs), anti-citrullinated protein antibodies (ACPA), and pro-inflammatory cytokines. B cells, in turn, mediate T cell activation through the expression of costimulatory molecules (1).

CD20 is a 33-37 kDa non-glycosylated protein that belongs to the membrane-spanning 4-domain A (MS4A) protein family (5), it is found in the majority of B cells starting from late pre-B cells, and its expression is lost in terminally differentiated plasmablasts and plasma cells (6, 7). B cell depletion by targeting CD20 using anti-CD20 monoclonal antibodies (mAbs) such as rituximab (RTX) has proved beneficial in RA (8, 9) and other autoimmune conditions, including multiple sclerosis (MS), neuromyelitis optica, and myasthenia gravis (10–13). The rationale for targeting anti-CD20 mAbs in RA was that removing autoantibody-producing or T cell-activating B cells would lead to clinical improvement (14, 15). However, although RTX reduces autoantibody titres, this effect did not explain its anti-inflammatory activity, which develops before the decrease in autoantibody titres is observed.

Multiple analyses have consistently demonstrated the expression of CD20 by a subset of CD3^+^ T cells (16–21). CD3^+^CD20^+^ T cells were found in the peripheral blood of RA patients and other human autoimmune syndromes, such as multiple sclerosis (MS), Primary Sjogren’s syndrome, and Psoriasis (17–19, 21). This CD20 expressing T cell subset exhibited pro-inflammatory features such as IFNγ, IL-1β, IL-17, IL-2, IL-8, transforming growth factor β (TGF-β) and TNFα, and C-C chemokine receptor type 2 (CCR2), CCR5, CCR6; these cytokines and chemokine receptors play crucial roles in homeostasis as well as pathologic immune contribution in various diseases, including RA (17–19). Like CD20^+^ B cells, CD3^+^CD20^+^ T cells were eliminated with RTX treatment (10, 17, 19–21). Hence, the therapeutic effect of anti-CD20 mAbs has also been attributed to the depletion of this highly inflammatory CD20^+^ T cell subset (21).

Animal models of RA have provided important insights into basic pathogenic mechanisms of chronic inflammatory arthritis and autoimmune disease in general. Collagen-Induced Arthritis (CIA) is a well established experimental model of RA to study both pathogenesis and its underlying immunological basis. It shares several critical characteristics of the disease pathogenesis with RA, including CD4^+^ T cell-mediated inflammation and extensive cartilage and bone damage, resulting in joint deformities (22, 23). Susceptibility to CIA is related to the murine MHC class II molecule H-2q whose peptide-binding pocket has a similar primary structure to the shared epitope of RA-associated HLA-DR molecules (24, 25).

The prevalence and contribution of CD20^+^ T cells in RA have largely been examined in the peripheral blood; however, the inflammatory role of these cells at the site of inflammation is unclear. In this study, we used the CIA model to investigate the phenotype and functional relevance of CD20 expressing T cells in the lymph nodes and arthritic joints of CIA mice using multicolor flow cytometry and immunohistochemistry techniques (22, 23). We report that draining lymph nodes and inflamed tissue of DBA1/J mice habour CD3^+^CD4^+^CD20^+^ and CD3^+^CD8^+^CD20^+^ T cells. These cell populations are expanded in mice undergoing CIA, and they display a pro-inflammatory phenotype and exhibit features associated with promoting inflammatory B cell responses. Our findings are suggestive that CD20^+^ T cells might contribute to pro-inflammatory processes in CIA.

## Materials and Methods

### Mice and induction of CIA

DBA1/J mice (8-10-week-old) were purchased from Envigo and were maintained under 12 h light/dark cycles and standard temperature (20-25 °C) in the University of Glasgow Biological Services Units in accordance with the Home Office UK Licenses P8C60C865, I675F0C46, and ID5D5F18C, and the Ethics Review Board of the University of Glasgow. Collagen-Induced Arthritis (CIA) was induced with bovine type II Collagen (MD Biosciences, 100 mg), injected intradermally (day 0) in Freund’s complete adjuvant (CFA). On day 21, mice received 100 mg of collagen in PBS intraperitoneally. Disease scores were measured every 48 h on a scale from 0 to 4 for each limb. An overall score of 10 or more was considered an experimental endpoint, at which point mice were immediately euthanized. For subsequent experiments, CIA mice were divided into arthritic and non-sick, showing clinical signs (Scores >4) or lacking clinical signs, respectively. Naïve non-immunised mice were used as control.

### Flow cytometric analysis

Popliteal and axillary draining lymph nodes were removed and ground through a 40 μm nylon mesh. Briefly, after washing, cells were either left unstimulated or activated for 4 h with phorbol 12-myristate 13-acetate (50 ng/ml; Sigma-Aldrich) and ionomycin (500 ng/ml; Sigma-Aldrich) in the presence of brefeldin A (1 μg/ml; Merck). The cells were washed and then stained with Fixable Viability Dye (eBioscience) for dead cell discrimination. Following a washing step, cells were labeled with fluorochrome antibodies specific for CD3-BUV395, CD4-BV510, CD8-BUV805, CD19-AF700, CXCR5-PerCp-Cy5.5, PD-1-PE-Dazzle (BD Biosciences), and CD20-FITC (Biolegend). For intracellular staining, the labeling was performed using antibodies specific for GM-CSF-BV421, TNF-α-APC, IL-17A-PE-Dazzle, and IFN-γ-BV711 (Biolegend). Appropriate isotype controls were used. Data were acquired using an LSR-Fortessa cell analyser (BD Biosciences). Flow cytometry datasets were analysed using FlowJo (version 10.7, Tree Star, Ashland, OR, USA).

Expression of CD20, CXCR5, and PD-1 was investigated in cells isolated from joints of non-arthritic and arthritic mice. Flow cytometry data were analysed using opt-SNE, an automated optimised algorithm for embedding cytometry data that improves visualisation and analysis of large datasets (26). Briefly, flow cytometry files corresponding to non-arthritic and arthritic mice were uploaded onto the premium version of Cytobank (Cytobank, Inc; www.cytobank.org). Events were first gated for singlets for exclusion of debris, and then lymphocytes were depicted based on the FSCA vs. SSCA gate. T cells were identified within the lymphocyte population based on CD3 expression, and then CD3^+^CD4^+^ and CD3^+^CD8^+^ T cells were identified within the CD3^+^ gate. CD3^+^CD4^+^ and CD3^+^CD8^+^ T cells were subjected to Opt-SNE based on CD20, CXCR5, and PD-1. The default settings of opt-SNE were used unless otherwise noted.

### Immunofluorescent staining of the lymph node and inflamed joint tissue sections

Lymph nodes were collected from mice, fixed in 10% neutral buffered formalin (Sigma, HT501320-9.5L), and moved into OCT embedded frozen blocks. Whole legs were harvested from animals, fixed in 10% neutral buffered formalin for 24 h, and transferred to 70% ETOH. Specimens were decalcified in EDTA and embedded in paraffin blocks. Tissue sections (7 μm) were deparaffinized in xylene and dehydrated in ethanol, and the antigen was retrieved in Citrate buffer (TCS Biosciences, catalog number: HDS05-100) at 95°C (pH 6.0) for 20 min. The sections were washed by PBS-T (PBS +0.05% Tween 20) for 5 min twice. Samples were stained with a rabbit anti-mouse CD20 antibody (1:100) (Cell Signaling, catalog number: 98708S) and a biotinylated hamster anti-mouse CD3 antibody (STEMCELL, catalog number: 60015BT.1; 1:100) at 4°C overnight. Sections were washed with PBS-T at least 3 times. Streptavidin-Alexa 647 (Invitrogen, catalog number: S32357) and Alexa 555-goat anti-rabbit secondary antibody (Invitrogen, catalog number: A21428; 1:400) were applied in PBS at room temperature for 60 min. Slides were rinsed with PBS and mounted with mounting media with DAPI (Invitrogen, catalog number: S36964). Images were obtained using an *LSM 880* confocal microscope and analyzed with Image J software.

### Statistical analyses

Data were analysed using the non-parametric Mann-Whitney *U* test with GraphPad Prism version 9.1.0. The summary data are presented as mean ± standard error of the mean (SEM) unless otherwise indicated. Sample sizes for all shown data can be found in the figure legends. *P* values less than 0.05 were considered significant.

## Results

### Increased frequencies of CD20^+^ T cells in the lymph nodes of arthritic mice

To identify CD20 expressing T cells and investigate potential implications in joint inflammatory disease, we isolated cells from popliteal and axillary draining lymph nodes of DBA1/J mice grouped as naïve (control), CIA mice that developed visible inflammation (arthritic), and those with no clinical signs despite being immunized (non-sick mice), which served as an additional control group. We performed multicolor flow cytometry for CD3, CD4, CD8, CD20, and CD19. Briefly, analyses were confined to live lymphocytes, and T cells were identified within the lymphocyte population based on CD3 expression, and then CD4^+^ and CD8^+^ T cells were identified within the CD3^+^ gate **(Figure 1A)**. CD20 expression was evaluated on CD3^+^CD4^+^ and CD3^+^CD8^+^ T cells and identified a proportion of CD4^+^CD20^+^ and CD8^+^CD20^+^ T cells in naïve, non-sick and CIA mice **(Figure 1B and C)**. We confirmed staining specificity using an appropriate isotype control, which did not show non-specific expression. Notably, CIA mice harbored higher proportions of CD4^+^CD20^+^ T cells than in naïve or non-sick mice **(Figure 1D).**

**Figure 1.**
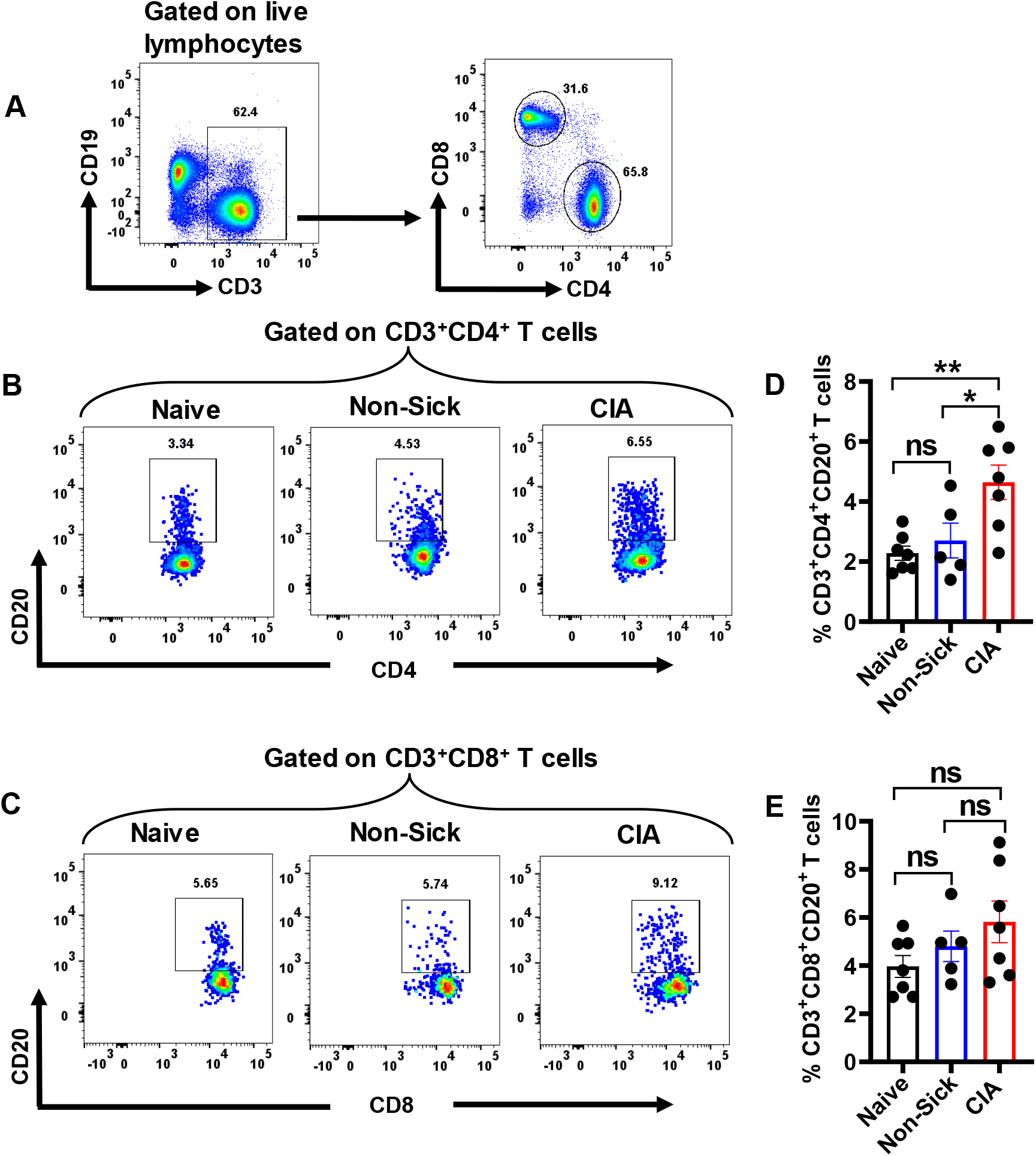
Increased frequencies of CD20^+^ T cells in the lymph nodes of CIA mice. (**A**) Flow cytometry dot-plots depicting approach used to gate on CD3^+^, CD4^+^, and CD8^+^ T cells. (**B and C**) Dot-plots showing representative Naïve, Non-Sick, and CIA mice, depicting CD20 expression by CD4^+^ and CD8^+^ T cells. (**D and E**) Percentages of CD3^+^CD4^+^CD20^+^ and CD3^+^CD8^+^CD20^+^ T cells in Naïve (n=7), Non-Sick (n=5) and CIA (n=7) mice. *p<0.05, **p<0.01. ns, not significant.

In contrast, although the percentage of CD20 expressing cells was higher within CD8^+^ T cells than in CD4^+^ T cells, the proportion of CD8^+^CD20^+^ T cells did not differ between naïve mice, non-sick mice, or CIA mice **(Figure 1E)**. These data suggested that the expansion of CD20^+^ T cells was associated with inflammatory arthritis and not systemic inflammation, as non-sick mice were treated with CFA but were similar to naïve controls.

### CD20^+^ T cells in CIA mice exhibit increased pro-inflammatory cytokine responses

To evaluate the functional properties of CD20^+^ T cells, we stimulated the lymph node cells with PMA/ionomycin and measured the cytokine production by CD4^+^CD20^+^ and CD8^+^CD20^+^ T cells using flow cytometry. We focused on GM-CSF, TNF-α, IL-17, and IFN-γ cytokines because of their role in promoting synovial inflammation, cartilage destruction, and bone erosion. We found that CD4^+^CD20^+^ T cells of CIA mice expressed higher percentages of GM-CSF, TNF-α, IL-17, and IFN-γ compared to naïve and non-sick mice **(Figure 2A-D)**. Within the CD8^+^CD20^+^ T cell population, CIA mice harbored increased frequencies of GM-CSF and IFN-γ expressing cells compared with naïve and non-sick mice, while no difference in TNF-α or IL-17 expression was observed between CIA, naïve and non-sick mice **(Figure 2E-H)**.

**Figure 2.**
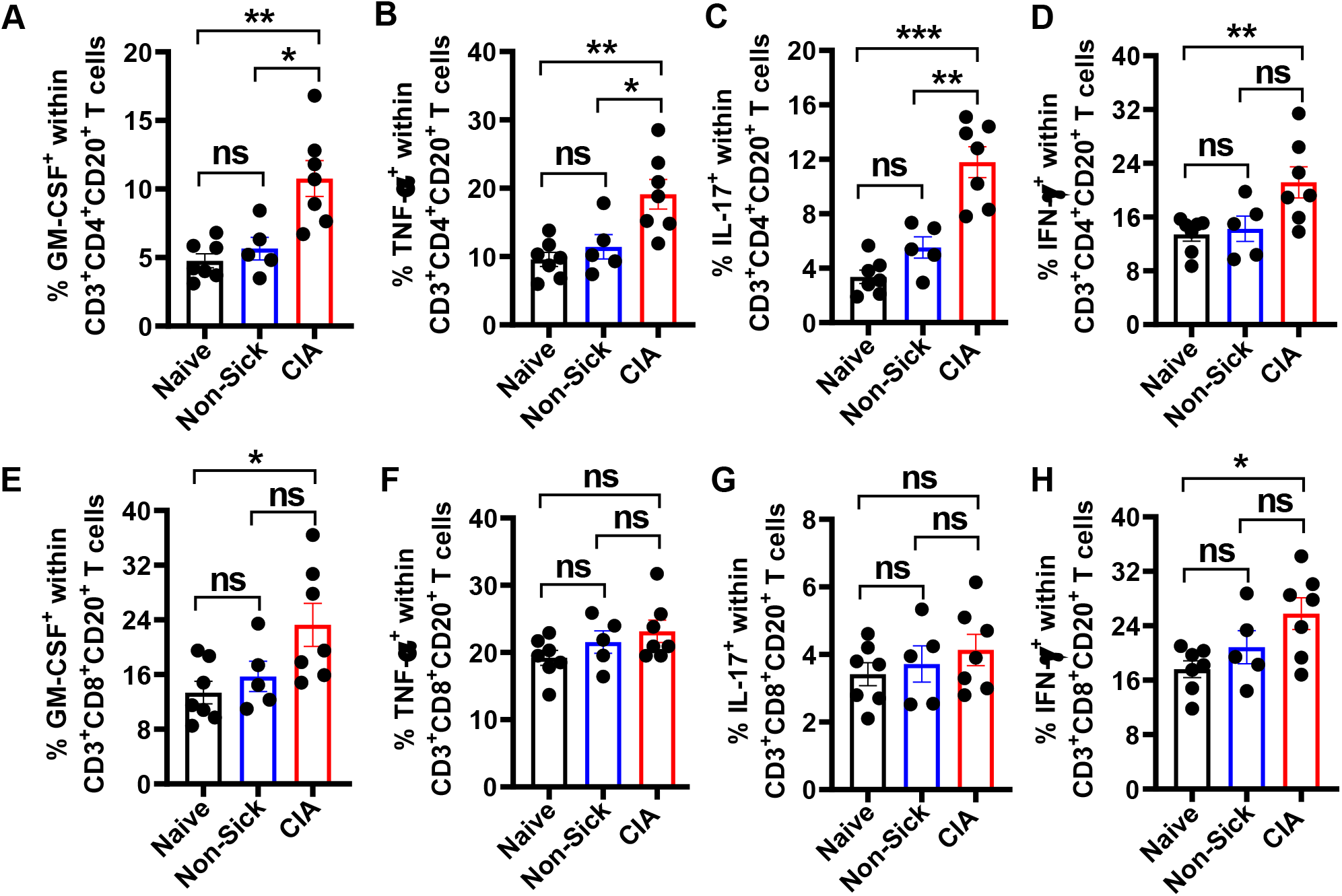
Increased proinflammatory cytokine responses of CD20 expressing T cells in CIA mice. The frequency of GM-CSF (**A**), TNF-α (**B**), IL-17 (**C**), and IFN-γ (**D**) producing CD3^+^CD4^+^CD20^+^ T cells in Naïve, Non-Sick, and CIA mice. (**E-H**) Percentages of GM-CSF, TNF-α, IL-17 and IFN-γ expression by CD3^+^CD8^+^CD20^+^ T cells in Naïve, Non-Sick and CIA mice. Naïve, n=7; Non-Sick, n=5; CIA, n=7. *p<0.05, **p<0.01; ***p<0.001. ns, not significant.

### CD3^+^CD20^+^ T cells are present at sites of inflammation in the arthritic joints

Next, we sought to determine the presence of CD20 expressing T cells in the inflamed regions. We stained the lymph node and joint sections with anti-CD3 and anti-CD20 antibodies and DAPI as a nuclear counterstain. The lymph node tissues contained extensive recruitment of CD3^+^ T cells. Notably, co-expression of CD3 and CD20 was seen in the inflamed areas **(Figure 3A).** Immunostaining also revealed co-expression of CD3 and CD20 in the arthritic joints **(Figure 3B).**

**Figure 3.**
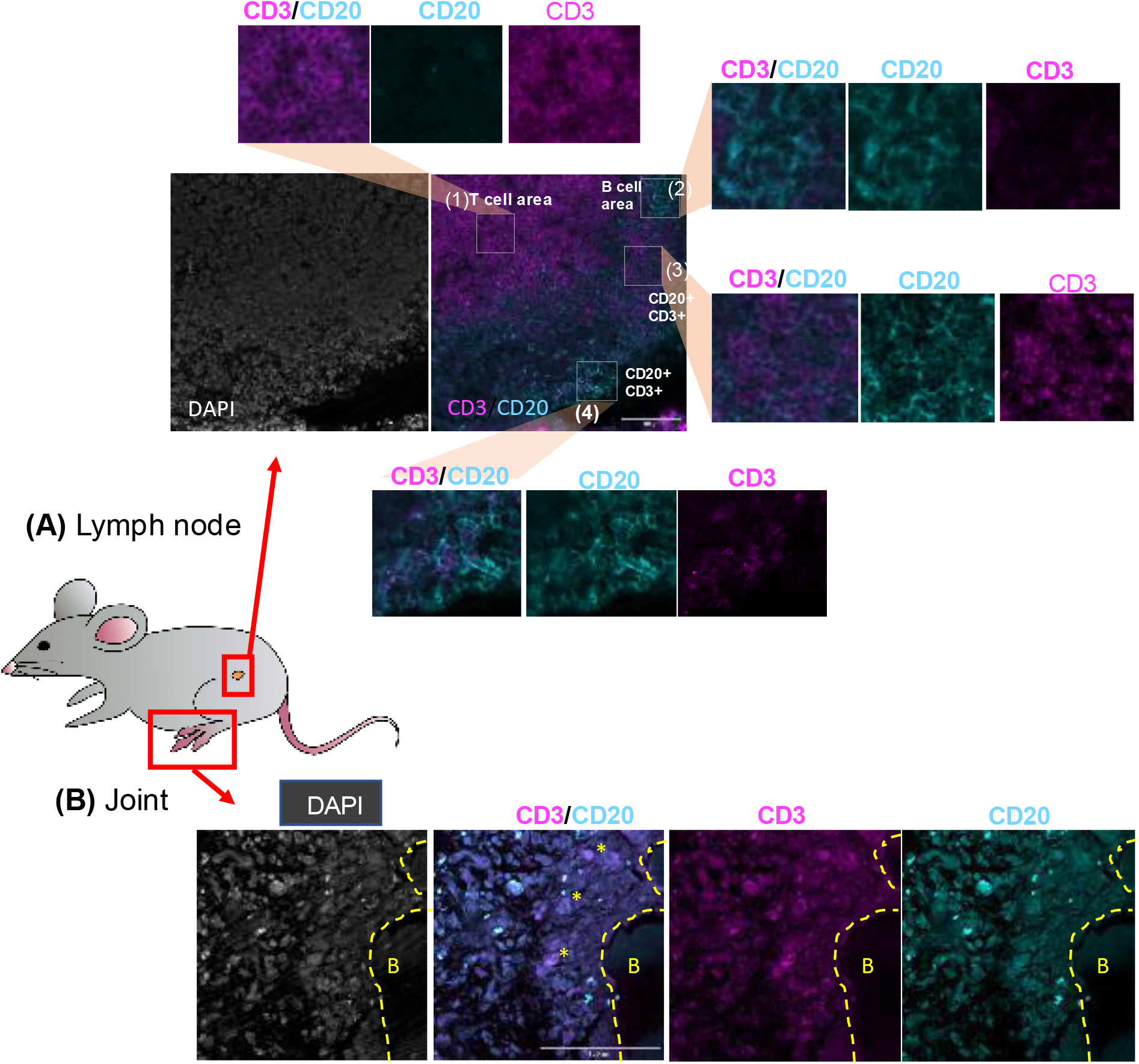
Expression of CD20 and CD3 in lymph nodes and joint tissue of CIA mice. Lymph nodes (**A**) and joint tissue (**B**) from mice undergoing CIA were collected and fixed in formaldehyde. Sections were stained with specific anti-CD3 (Magenta) and anti-CD20 (Cyan) antibodies. DAPI (grey) was used to stain nucleic acid as counterstaining. Scale bars: 100 μm. Superimposed yellow elements indicate as follows; [dotted lines]: bone limits, [asterisks]: synovial CD20^+^CD3^+^ cells. Images were acquired using a confocal microscope.

### CXCR5^+^PD-1^+^ and CXCR5^-^PD-1^+^ T cells are enriched within CD3^+^CD4^+^CD20^+^ and CD3^+^CD8^+^CD20^+^ T cells, particularly in lymph nodes of CIA mice

CD20^+^ T cells have been suggested to arise due to intercellular membrane exchange during intimate T cell / B cell interaction. In line with this, we wondered whether CD20^+^ T cells exhibit features enabling B cell help. Therefore, we assessed co-expression of CD20, CXCR5, and PD-1 on CD3^+^CD4^+^ and CD3^+^CD8^+^ T cells. First, we found that CD3^+^CD4^+^ T cells of CIA mice expressed increased frequencies of CXCR5^+^PD-1^+^ T follicular helper (T_FH_) cells compared to naïve and non-sick mice **(Figure 4A and B)**. Also, compared to naïve and non-sick mice, CD3^+^CD4^+^ T cells of CIA mice harbored higher frequencies of CXCR5^-^PD-1^+^ T cells **(Figure 4A and C)**, a subset that has been defined as peripheral helper T (T_PH_) cells (27). Notably, CXCR5^+^PD-1^+^ T_FH_ and CXCR5^-^PD-1^+^ T_PH_ were enriched within the CD3^+^CD4^+^CD20^+^ T cell population, and increased frequencies were found in CIA mice compared to naïve or non-sick mice **(Figure 4D, E, and F)**. In contrast, CD3^+^CD4^+^CD20^-^ T cells harbored lower frequencies of CXCR5^+^PD-1^+^ T_FH_ or CXCR5^-^PD-1^+^ T_PH_, and no differences were observed between naïve, non-sick, and CIA mice **(Figure 4G, H and I).**

**Figure 4.**
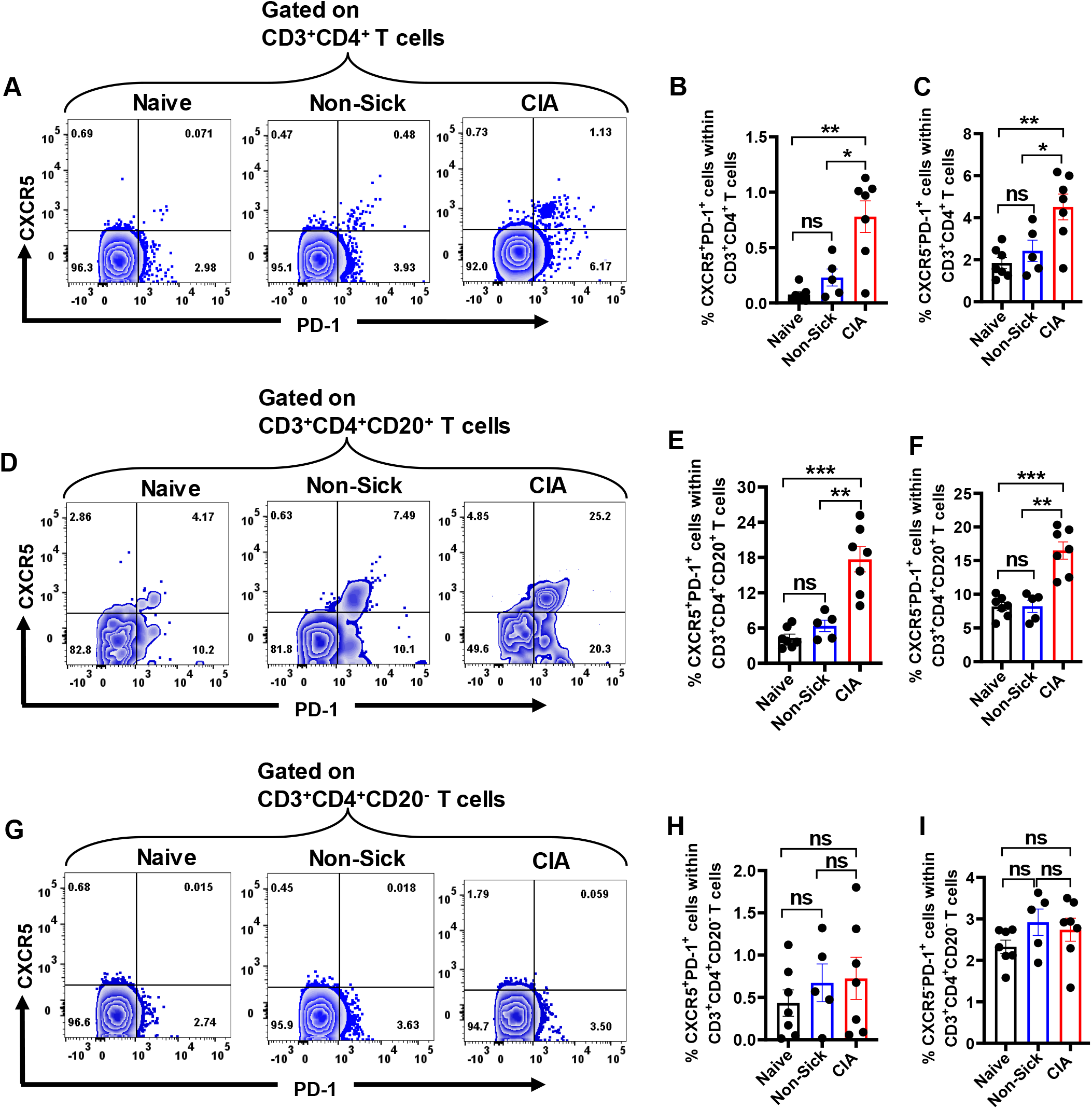
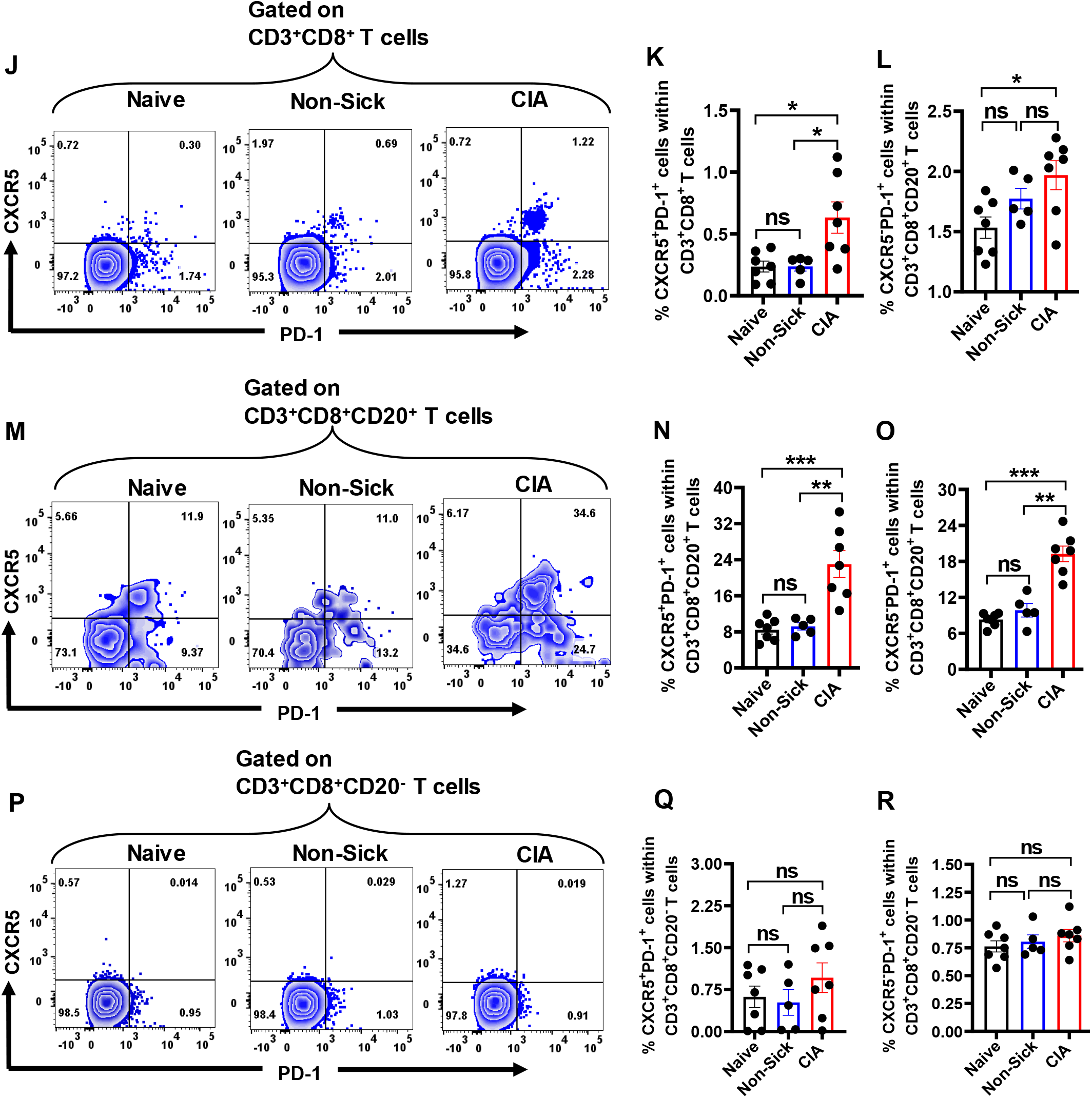
CXCR5^+^PD-1^+^ and CXCR5^-^PD-1^+^ T cells are enriched within CD3^+^CD4^+^CD20^+^ and CD3^+^CD4^+^CD20^+^ T cells, particularly in CIA mice. (**A**) Flow cytometry dot-plots depicting CXCR5 and PD-1 expression by total CD4^+^ T cells. (**B, C**) Percentages of CXCR5^+^PD-1^+^ and CXCR5^-^PD-1^+^ T cells by total CD3^+^CD4^+^ T cells. (**D**) Flow cytometry dot-plots depicting CXCR5 and PD-1 expression within CD3^+^CD4^+^CD20^+^ T cells in representative Naïve, Non-Sick, and CIA mice. (**E, F**) Percentages of CXCR5^+^PD-1^+^ and CXCR5^-^PD-1 ^+^ T cells among CD3^+^CD4^+^CD20^+^ T cells, respectively. (**G**) Flow cytometry dotplots depicting expression of CXCR5 and PD-1 within CD3^+^CD4^+^CD20^-^ T cells. (**H, I**) Percentages of CXCR5^+^PD-1^+^ and CXCR5^-^PD-1^+^ T cells among CD3^+^CD4^+^CD20-T cells, respectively. (**J**) Flow cytometry dot-plots depicting CXCR5 and PD-1 expression by total CD3^+^CD8^+^ T cells. (**K, L**) Percentages of CXCR5^+^PD-1^+^ and CXCR5^-^PD-1^+^ T cells within CD3^+^CD8^+^ T cells. (**M**) Flow cytometry dot-plots depicting expression of CXCR5 and PD-1 within CD3^+^CD8^+^CD20^+^ T cells. (**N, O**) Percentages of CXCR5^+^PD-1^+^ and CXCR5^-^PD-1^+^ T cells among CD3^+^CD8^+^CD20^+^ T cells, respectively. (**P**) Flow cytometry dot-plots depicting expression of CXCR5 and PD-1 within CD3^+^CD8^+^CD20^-^ T cells. (**Q, R**) Percentages of CXCR5^+^PD-1^+^ and CXCR5^-^PD-1^+^ T cells within CD3^+^CD8^+^CD20^-^ T cells. Naïve, n=7; Non-Sick, n=5; and CIA mice, n=7. *p<0.05, **p<0.01. ns, not significant.

The CXCR5^+^PD-1^+^ T_FH_ and CXCR5^-^PD-1^+^ T_PH_ subsets were also found on CD3^+^CD8^+^ T cells, and higher frequencies were detected in CIA mice than in naïve or non-sick mice **(Figure 4J, K and L)**. Compared to the frequencies of CXCR5^+^PD-1^+^ T_FH_ and CXCR5^-^PD-1^+^ T_PH_ within total CD3^+^CD8^+^ T cells, CXCR5^+^PD-1^+^ T_FH_ and CXCR5^-^PD-1^+^ T_PH_ were enriched within the CD3^+^CD8^+^CD20^+^ T cell population, and significantly higher levels were found in CIA mice than in naïve and non-sick mice **(Figure 4M, N, and O)**. Relatively, CD3^+^CD4^+^CD20^-^ T cells contained lower frequencies of CXCR5^+^PD-1^+^ T_FH_ or CXCR5^-^PD-1^+^ T_PH_, and no differences were observed between naïve, non-sick, and CIA mice **(Figure 4P, Q and R)**.

### Visualisation of CD20, CXCR5, and PD-1 co-expressing cells in CD3^+^CD4^+^ and CD3^+^CD8^+^ T cells in non-arthritic and arthritic paws

To examine whether T cells at the site of inflammation express CD20, CXCR5, and PD-1, the joint tissues of non-arthritic and arthritic mice were digested with collagenase II and the single-cell suspension was analyzed by flow cytometry. The flow cytometry data were explored using opt-SNE, a powerful optimization toolkit that subverts significant t-SNE limitations for use with cytometric datasets and thus enables novel data-driven findings in single-cell data (26). CD3^+^CD4^+^ and CD3^+^CD8^+^ T cells were subjected to opt-SNE based on CD20, CXCR5, and PD-1 expression. Opt-SNE helped visualize the co-expression of CD20, CXCR5, and PD-1 in CD4^+^ T cells of non-arthritic and arthritic mice (**Figure 5A**). The CD4^+^ T cell population was exported, and the mean fluorescence intensity (MFI) of CD20, CXCR5, and PD-1 was assessed using FlowJO software. The intensity of CD20 expression was higher in CD4^+^ T cells of arthritic mice than non-arthritic mice (**Figure 5B**; right and left panels). In contrast, the intensities of CXCR5 and PD-1 did not differ between arthritic and non-arthritic mice (**Figure 5C and D**; right and left panels, respectively).

**Figure 5.**
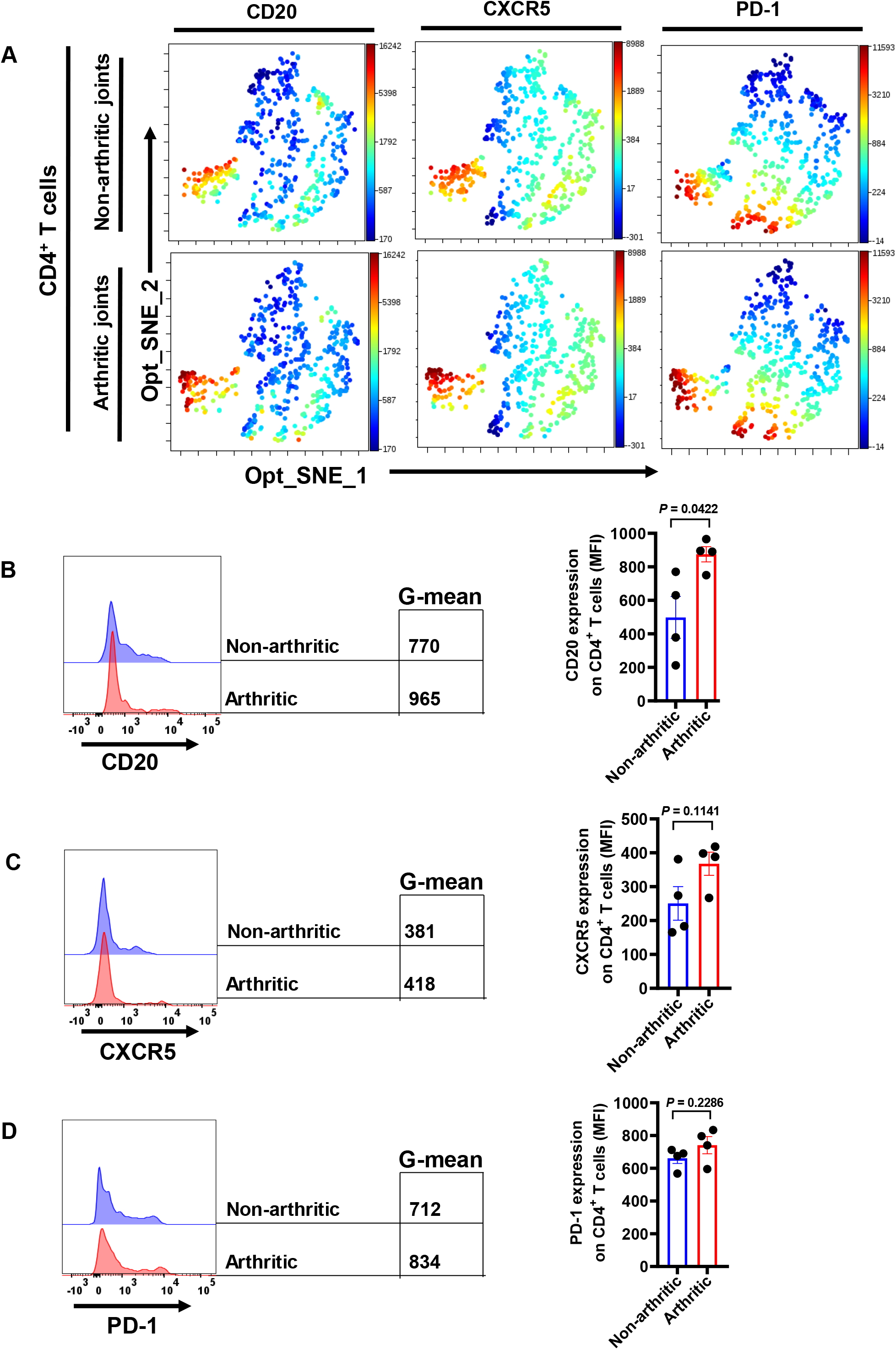
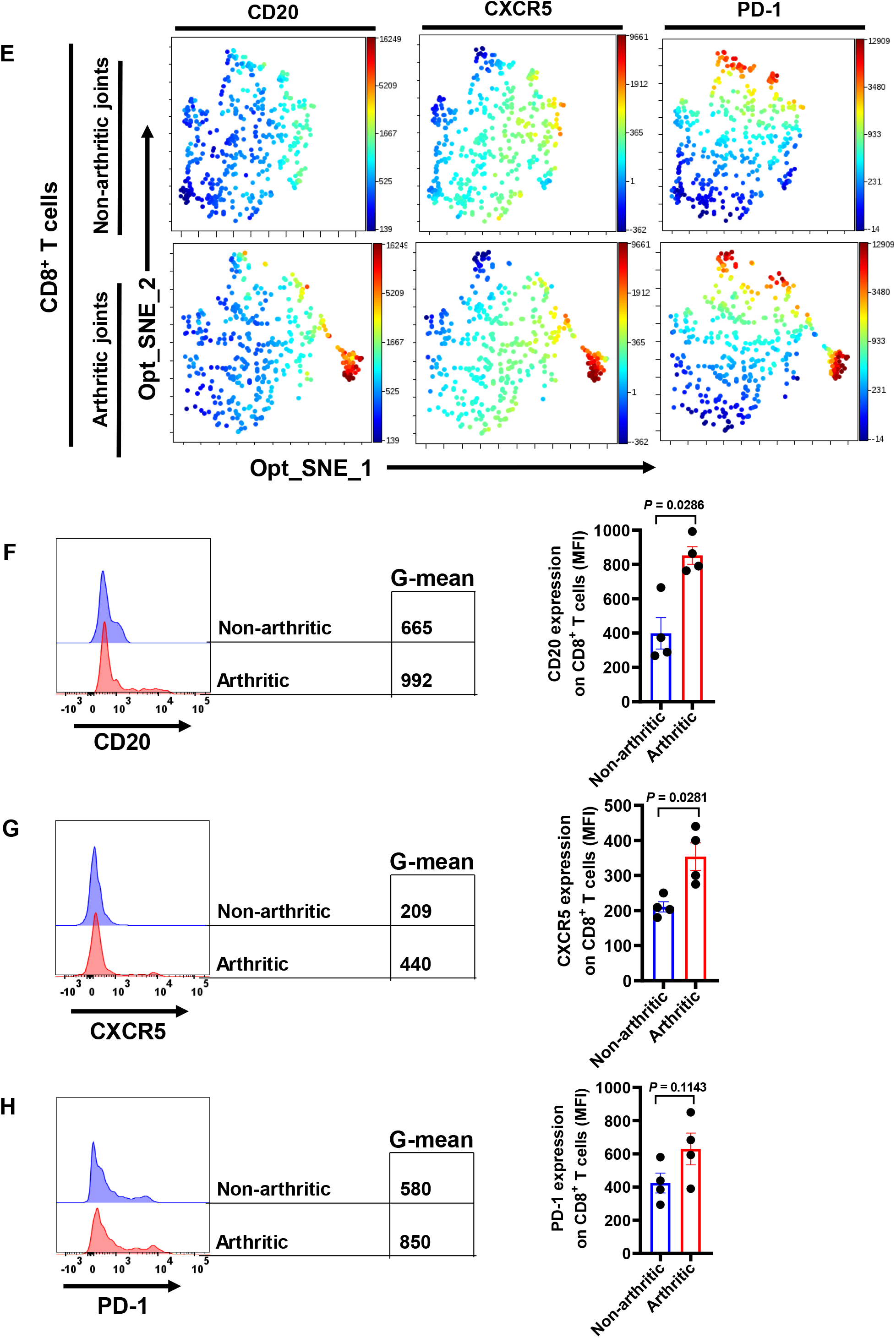
Visualisation of CD20, CXCR5, and PD-1 co-expressing cells in CD3^+^CD4^+^ and CD3^+^CD8^+^ T cells in non-arthritic and arthritic joints. Flow cytometry dataset files from single, live CD4^+^ or CD8^+^ T cells from non-arthritic and arthritic joints were used as input for the opt-SNE embedding using Cytobank. (**A**) CD20, CXCR5, and PD-1 expression by CD4^+^ T cells of non-arthritic and arthritic mice are depicted. (**B-D**) Mean fluorescence intensity (MFI) of CD20, CXCR5, and PD-1 and summary data are presented. (**E**) CD20, CXCR5, and PD-1 expression by CD8^+^ T cells of non-arthritic and arthritic mice are shown. (**F-H**) Mean fluorescence intensity (MFI) of CD20, CXCR5, and PD-1 in CD8^+^ T cells and summary data are presented. Non-arthritic mice, n=4; Arthritic mice; n=4.

Within CD3^+^CD8^+^ T cells, Opt-SNE demonstrated the co-expression of CD20, CXCR5, and PD-1 in non-arthritic and arthritic mice (**Figure 5E**). Analysis of the exported CD3^+^CD8^+^ T cell population showed that arthritic mice had higher CD20 and CXCR5 expression intensity than non-arthritic mice (**Figure 5F and G**; right and left panels, respectively). However, the intensity of PD-1 did not differ between arthritic and non-arthritic mice (**Figure 5H**; right and left panels). Together, these data confirm the expansion of CD20^+^ T cells in CIA mice and suggest an association with inflammatory responses.

## Discussion

This study provides evidence that CD20^+^ T cells are present in the lymph nodes and arthritic joints of CIA mice, and this CD20^+^ T cell pool comprises both CD4^+^ and CD8^+^ T cells. We show that the frequencies of CD3^+^CD4^+^CD20^+^ and CD3^+^CD8^+^CD20^+^ T cell populations are increased in mice with CIA, producing high levels of pro-inflammatory cytokines (GM-CSF, TNF-α, IL-17, and IFN-γ). Further, we show that CD20^+^ T cells are enriched with CXCR5^+^PD-1^+^ T_FH_ and CXCR5^-^PD-1^+^ T_PH_ cells, subsets that have been implicated in promoting inflammatory B cell responses (27, 28).

The presence of CD20^+^ T cells has been reported in the blood of healthy controls and autoimmune diseases (20, 21), including RA (18). In non-human primates, CD3^+^CD20^+^ T cells were found in the lymph nodes of monkeys (29). Although a CD20 homolog, MS4aB1, was detected on murine CD3^+^ T cells (30, 31), no murine CD3^+^CD20^+^ T cells have previously been reported. To our knowledge, this is the first report to show that CD20^+^ T cells are present in the lymph nodes and the joints of mice, and that this population is expanded in mice with severe joint inflammation. We used 8-10 weeks DBA1/ mice, the incidence of developing CIA in our model was 60-70% (scores >2). Therefore, we took advantage of the CIA model not showing 100% incidence; hence, we compared animals with a severe disease with animals that did not develop inflammation despite receiving CFA. In our evaluation of the functional role of CD3^+^CD20^+^ T cells in the development of autoimmune joint inflammation, we phenotyped these cells in naïve mice (controls), CIA mice that developed visible inflammation (arthritic) as well as those with no clinical signs of arthritis despite being immunised (non-sick mice). Hence, the non-sick mice served as an additional control group in addition to the naïve control mice. Comparatively, CIA mice harbored a higher percentage of CD4^+^CD20^+^ T cells than naïve and non-sick mice. Although the percentage of CD8^+^CD20^+^ T cells did not significantly differ between naïve, non-sick, and CIA mice, a higher frequency was observed in CIA mice compared to naïve and non-sick mice, and the proportion of CD20 expressing cells was higher within CD8^+^ than in CD4^+^ T cells.

The observation of increased production of IL-17, TNF-α, IFN-γ, and GM-CSF by CD4^+^CD20^+^ T cells, and excessive production of GM-CSF and IFN-γ by CD8^+^CD20^+^ T cells, particularly from arthritic mice, is consistent with the suggestion that CD20^+^ T cells might play a crucial role in pro-inflammatory processes in autoimmune conditions (17–21). Studies on human immune cells showed that peripheral blood of RA patients harbored high proportions of IL-17-producing CD20^+^ T cells compared to healthy individuals (17, 18). CD3^+^CD20^+^ T cells have also been found in the brain tissues of MS patients, exhibiting high proportions of IL-17 and IFN-γ producing cells in the peripheral blood (19, 20, 32). In Psoriasis patients, CD3^+^CD20^+^ T cells showed increased IL-17, TNF-α, and IL-21, while high proportions of IL-17 producing CD3^+^CD20^+^ T cells were reported in the blood of individuals with Primary Sjogren’s syndrome (33, 34). IL-17 plays a critical inflammatory role in the propagation of CIA and RA (35–37); it acts on the osteoclastogenesis supporting cells (38) and facilitates local inflammation by activating immune cells and recruiting them to the site of inflammation (39, 40). The expression of GM-CSF by CD4^+^CD20^+^ and CD8^+^CD20^+^ T cells further demonstrates their pathological role in CIA. Notably, GM-CSF has been implicated in the inflammatory context observed in many autoimmune diseases, such as RA and MS (RA) (41, 42), and recent data show that GM-CSF is an essential cytokine in RA development (43, 44). Since the potential pathological effects of CD3^+^CD20^+^ T cells occur in the inflamed regions, we examined the lymph node and joint sections for CD3^+^CD20^+^ T-cells. We observed extensive recruitment of CD3^+^ T cells co-expressing CD20 in the inflamed areas of the lymph nodes and the arthritic joints of CIA mice. Thus, our data associate CD3^+^CD20^+^ T cells with an inflammatory role in CIA.

We established that CD3^+^CD4^+^CD20^+^ T cells are enriched with CXCR5^+^PD-1^+^ T cells. CD4^+^ T_FH_ cells have been characterized by their expression of CXCR5 (45, 46) and programmed cell death-1 (PD-1) (47). T_FH_ cells provide help to antigen-specific B cells; this helper function is the major role of T_FH_ cells, a specialized subset of CD4^+^ T cells that localize to B-cell follicles and provide the cognate B-T cell interaction to facilitate the formation of germinal centers, promote the differentiation of germinal center B cells into memory B or plasma cells, and drive the development of high-affinity antibodies (48, 49). T_FH_ cells have been implicated in the development of autoimmune diseases, including in animal models where dysregulated T_FH_ function promoted autoantibody formation (50, 51). In humans, increased T_FH_ cell numbers were identified in patients with autoimmune diseases such as RA and systemic lupus erythematosus (SLE) (52–54).

T_PH_ cells have been implicated in interactions between T-B cell collaboration, located adjacent to B cells inside and outside synovial lymphoid aggregates in patients with RA, and have been shown to actively promote memory B cell differentiation into plasma cells (27). We confirmed the existence of CXCR5^-^PD-1^+^ T_PH_ cells within the CD3^+^CD4^+^CD20^+^ T cell subset in our model. Previous studies showed that CXCR5^-^PD-1^+^CD4^+^ T cells generate ectopic lymphoid structures in the RA synovium (55, 56). The enrichment of T_FH_ and T_PH_ cells within CD4^+^CD20^+^ T cells suggests that these cells may be more efficient at supporting B cell responses than CD20^-^ T cells. However, further experiments are required to confirm this hypothesis. The observation of increased production of pro-inflammatory cytokines such as IL-17 by CD4^+^CD20^+^ T cells supports this suggestion. Moreover, it has been reported that Th17 cells had a higher ability to support antibody production and persistence in the B cell follicle and were in close proximity to antigen-specific B cells after antigen challenge (57).

Comparatively, less little attention has been paid to the role of CD8^+^ T cells than CD4^+^ T cells in RA or CIA; however, CD8^+^ T cells comprise about 40% of all T cells in RA synovium, and the abundance of these cells in the synovium and peripheral blood is associated with disease activity (58). Recent reports have highlighted the ability of CD8^+^ T cells to function as T_FH_ cells in the germinal center (GC), promoting B cell differentiation and antibody isotype class switching (59, 60). In RA synovial ectopic follicles, CD8^+^ T cells make up most of the infiltrating T cells (61) and are required to form ectopic GC in rheumatoid synovitis (60). CD8^+^ T cells expressing CXCR5 have been shown to develop under inflammatory conditions (62, 63), and CXCR5^+^ CD8 T cells are localized in the B cell follicle in the human tonsil were shown to support B cell survival in ex vivo culture (64). Further, increased proportions of the CXCR5^+^PD-1^+^CD8^+^ T cell subset have been found in the follicular B cell zones, and these CXCR5^+^PD-1^+^CD8^+^ T cells promoted B cells to produce autoantibodies in the absence of CD4^+^ T cells (28). Notably, CXCR5^+^PD-1^+^CD8^+^ T cells express CXCR5, a critical feature expressed on B cells and CD4^+^ T_FH_ cells, and directs cells to migrate into the B-cell follicles of the spleen and lymph nodes (27, 28, 59). Our observation that CD3^+^CD8^+^CD20^+^ T cells are enriched with CXCR5^+^PD-1^+^ T_FH_ cells suggests that, in addition to directly causing inflammation, the CD3^+^CD8^+^CD20^+^ T cells may cause additional pathology by promoting inflammatory B cell responses in CIA. Moreover, a recent report showed that a selective Cxcr5 deficiency in T cells (T-CXCR5^-/-^) hampered GC formation, weakened antibody response to CII upon CIA induction, and decreased serum levels of proinflammatory cytokines. Further, T-CXCR5^-^ mice did not develop arthritic paws (65).

A recent report showed that PD-1^+^CD8^+^ T cells also produce increased amounts of IL-21 that allow B cell differentiation; these cells were abundant in RA patients’ peripheral blood and synovial fluid. Of note, the PD-1^+^CD8^+^ T cell subset expressed CCR2 but not CXCR5 (66). Our data show that CD8^+^CD20^+^ T cells are enriched with CXCR5^-^PD-1^+^ T cells, and the levels were higher in CIA mice compared to naïve and non-sick mice, implicating CD8^+^CD20^+^ T cells in CIA induction and pathology.

In conclusion, we have demonstrated the presence of pro-inflammatory CD4^+^CD20^+^ and CD8^+^CD20^+^ T cells in the lymph nodes and arthritis paws of CIA mice. We have shown that the CD4^+^CD20^+^ T cell population is enriched with T_FH_ and T_PH_; CD8^+^CD20^+^ T cells also harbor subsets with similar features to T_FH_ and T_PH_. In agreement with suggestions in RA (66), our data suggest that CXCR5^+^PD-1^+^CD8^+^ T cells CXCR5^-^PD-1^+^CD8^+^ T cells in concert with T_FH_ and T_PH_ play an essential role in augmenting pathology in CIA. The capacity of CD4^+^CD20^+^ T cells to induce the expansion of cognate B cells and the production of class-switched antibodies warrants investigation.

## Notes

### Competing Interest Statement

The authors have declared no competing interest.

